# Functional correlates of immediate early gene expression in mouse visual cortex

**DOI:** 10.1101/2020.11.12.379909

**Authors:** David Mahringer, Pawel Zmarz, Hiroyuki Okuno, Haruhiko Bito, Georg B. Keller

## Abstract

During visual development, response properties of layer 2/3 neurons in visual cortex are shaped by experience. Both visual and visuomotor experience are necessary to coordinate the integration of bottom-up visual input and top-down motor-related input. Whether visual and visuomotor experience engage different plasticity mechanisms, possibly associated with the two separate input pathways, is still unclear. To begin addressing this, we measured the expression level of three different immediate early genes (IEG) (c-fos, egr1 or Arc) and neuronal activity in layer 2/3 neurons of visual cortex before and after a mouse’s first visual exposure in life, and subsequent visuomotor learning. We found that expression levels of all three IEGs correlated positively with neuronal activity, but that first visual and first visuomotor exposure resulted in differential changes in IEG expression patterns. In addition, IEG expression levels differed depending on whether neurons exhibited primarily visually driven or motor-related activity. Neurons with strong motor-related activity preferentially expressed EGR1, while neurons that developed strong visually driven activity preferentially expressed Arc. Our findings are consistent with the interpretation that bottom-up visual input and top-down motor-related input are associated with different IEG expression patterns and hence possibly also with different plasticity pathways.

## Introduction

During first visuomotor exposure in life, experience with coupling between movement and visual feedback is thought to coordination inputs onto layer 2/3 neurons in primary visual cortex (V1) such that individual neurons receive balanced and opposing top-down motor-related and bottom-up visual input (Attinger et al., 2017; Jordan and Keller, 2020; Leinweber et al., 2017). While visual input without visuomotor coupling is sufficient to establish normal visual responses in layer 2/3 neurons, the emergence of visuomotor mismatch responses is contingent on experience with visuomotor coupling (Attinger et al., 2017) and relies on NMDA receptor dependent signaling in the local V1 circuit (Widmer et al., 2022). How top-down and bottom-up inputs are coordinated during visual and visuomotor experience and whether plasticity mechanisms in the bottom-up visual driven pathway are the same as those engaged in the top-down motor-related input is still unclear. Here we set out to test for changes in expression levels of immediate early gene (IEG) products in functionally identified neurons during first exposure to visual input and first exposure to normal visuomotor coupling using concurrent measurement of neuronal activity and IEG expression levels in layer 2/3 of mouse visual cortex.

IEG products play a critical role in synaptic and neuronal plasticity during learning (Chowdhury et al., 2006; Fleischmann et al., 2003; Gandolfi et al., 2017; Jones et al., 2001; Messaoudi et al., 2007; Rial Verde et al., 2006; Shepherd and Bear, 2011; Shepherd et al., 2006; Tzingounis and Nicoll, 2006; Vazdarjanova et al., 2006; Veyrac et al., 2014; Waung et al., 2008) and are necessary for long-term memory consolidation (Bozon et al., 2003; Fleischmann et al., 2003; Guzowski, 2002; Guzowski and McGaugh, 1997; Guzowski et al., 2000; Jones et al., 2001; Ploski et al., 2008; Yasoshima et al., 2006). Ever since the discovery that the expression of the transcription factor c-Fos can be induced by electrical or chemical stimulation in neurons (Greenberg and Ziff, 1984), the expression of IEGs has been used as a marker for neuronal activity (Bullitt, 1990; Guzowski et al., 1999; Jarvis et al., 2000; Knapska and Kaczmarek, 2004; Minatohara et al., 2015; Morgan et al., 1987; Ramírez-Amaya et al., 2005; Reijmers et al., 2007; Wang et al., 2021). Based on the discovery that certain forms of episodic memory can be reactivated by artificially activating an ensemble of neurons characterized by high IEG expression levels during memory acquisition (Denny et al., 2014; Garner et al., 2012; Liu et al., 2012; Ramirez et al., 2013), it has been speculated that IEG expression is related not simply to neuronal activity *per se*, but to the induction of activity-dependent plasticity (Holtmaat and Caroni, 2016; Josselyn et al., 2015; Kaplan et al., 1996). Assuming IEG expression is indeed related to the induction of neuronal plasticity, it is conceivable that different IEGs are preferentially involved in plasticity of different synapse types or input pathways. c-Fos and EGR1 expression levels in visual cortex, for example, are differentially regulated by visual experience and exhibit a differential dependence on neuromodulatory input (Yamada et al., 1999). Consistent with a pathway-specific role of Arc and EGR1 in visual cortex, it has been shown that Arc is necessary for different forms of plasticity of bottom-up visual input, including ocular dominance plasticity (Gao et al., 2010; Jenks et al., 2017; McCurry et al., 2010; Wang et al., 2006), while a knockout of *egr1* has been shown to leave ocular dominance plasticity unaffected (Mataga et al., 2001), and EGR1 expression levels have been shown to be modulated in a context-specific manner, primarily in a subset of superficial layer 2/3 neurons (Xie et al., 2014). It is still unclear however, whether IEG expression is differentially regulated by changes in bottom-up and top-down input and if there is preferential expression of different IEGs in neurons that are predominantly excited by top-down motor-related input as compared to neurons that are predominantly excited by bottom-up visual input.

## Results

To measure both IEG expression levels and neuronal activity chronically, we used a combination of transgenic mice that express GFP under the control of an IEG promoter and viral delivery of a red variant of a genetically encoded calcium indicator. We did this for three different IEGs (*c-fos*, *egr1*, and *Arc*), in three groups of mice separately. EGFP-Arc and c-Fos-GFP mice are transgenic mice that express a fusion protein of Arc or c-Fos, and GFP downstream of either an *Arc* or a *c-fos* promoter, respectively (Barth et al., 2004; Okuno et al., 2012), while the EGR1-GFP mouse expresses GFP under an *egr1* promoter (Xie et al., 2014). Although there are a number of caveats to using GFP levels in these mouse lines as a proxy for IEG expression levels (see Discussion), there is a strong overlap between post-mortem antibody staining for the respective IEG and GFP expression in all three mouse lines (Barth et al., 2004; Okuno et al., 2012; Xie et al., 2014; Yassin et al., 2010). Throughout the manuscript, we will use IEG expression to mean GFP expression levels in these mice. To measure calcium activity, we used an AAV2/1-Ef1a-jRGECO1a viral vector to express the genetically encoded red calcium indicator jRGECO1a (Dana et al., 2016). This biased our recordings to excitatory neurons, as in the first few weeks after the injection, the Ef1a promoter restricts expression mainly to excitatory neurons (Attinger et al., 2017).

To quantify the correlation between neuronal activity and IEG expression of individual neurons in layer 2/3 of visual cortex in adult mice, we first used a paradigm of dark adaptation and subsequent brief visual exposure (**Figure 1A**). We did this in three groups of adult mice separately (4 EGFP-Arc mice, 4 c-Fos-GFP mice, and 4 EGR1-GFP mice, between 100 and 291 days old). We dark-adapted all three groups of mice for 24 hours and subsequently head-fixed them, while still in complete darkness, under a two-photon microscope on a spherical treadmill (**Figure 1A**). We then measured calcium activity and IEG levels every 15 minutes for six hours (**Figures 1B–1D**; see Methods). Between the first and second measurement, mice were exposed to visual input for 15 minutes. This paradigm, which is a combination of light exposure and exposure of the mouse to head-fixation, resulted in transient and modest increases in Arc and EGR1 expression levels, and a decrease in c-Fos expression levels (**Figure S1A**). Given that mean neuronal activity levels in V1 are rapidly stabilized across light-dark transitions (Hengen et al., 2016), and only a prolonged dark adaption (of 60 hours) results in a substantial increase of neuronal activity upon light exposure (Torrado Pacheco et al., 2019), the absence of a response in mean IEG levels is perhaps not surprising. We then computed the correlation between average neuronal activity and IEG expression levels for each neuron as a function of time between neuronal activity measurement and IEG expression measurement (**Figures 1E–1G**). Correlation peaked at a time lag of approximately 3.5 h ± 0.5 h (mean ± SEM) between neuronal activity measurement and IEG measurement for Arc and c-Fos, and was positive but relatively stable in a window from −2 hours to +3 hours for EGR1, consistent with previous results (Wang et al., 2021) (Arc: 1382 neurons, c-Fos: 1070 neurons, EGR1: 1319 neurons; **Figures 1E–1G**). At peak, the correlation between neuronal activity and IEG expression was highest for c-Fos, intermediate for Arc, and lowest for EGR1 (**Figures 1H–1J**; correlation coefficients for c-Fos: 0.39 ± 0.07, Arc: 0.26 ± 0.05, EGR1: 0.21 ± 0.03, mean ± SEM; comparisons between c-Fos vs. Arc: p < 3 × 10^-4^, Arc vs. EGR1: p = 0.0188, c-Fos vs. EGR1: p < 10^-8^; 4 mice per group, t-test with bootstrapping, see Methods). The positive correlation and the time lag of the correlation peak would be consistent with the idea that neuronal activity induces IEG expression, but the fact that correlations with mean activity were relatively weak could mean that it is specific patterns or types of activity that induce IEG expression. It is often assumed that IEG expression is also a correlate of neuronal plasticity (Holtmaat and Caroni, 2016; Josselyn et al., 2015; Kaplan et al., 1996). Given that certain forms of neuronal plasticity are associated with bursts of activity, we first tested whether maximum activity was a better predictor of IEG expression levels than mean activity. Indeed, we found that the correlation with maximum neuronal activity was higher than the correlation with mean activity for all three IEGs, but only significantly so for Arc and EGR1 (**Figure S1B**). Thus, while there is a weak but positive correlation between calcium activity and IEG expression for all three IEGs, it is possible that IEG expression is more directly related to functional plasticity.

**Figure 1.**
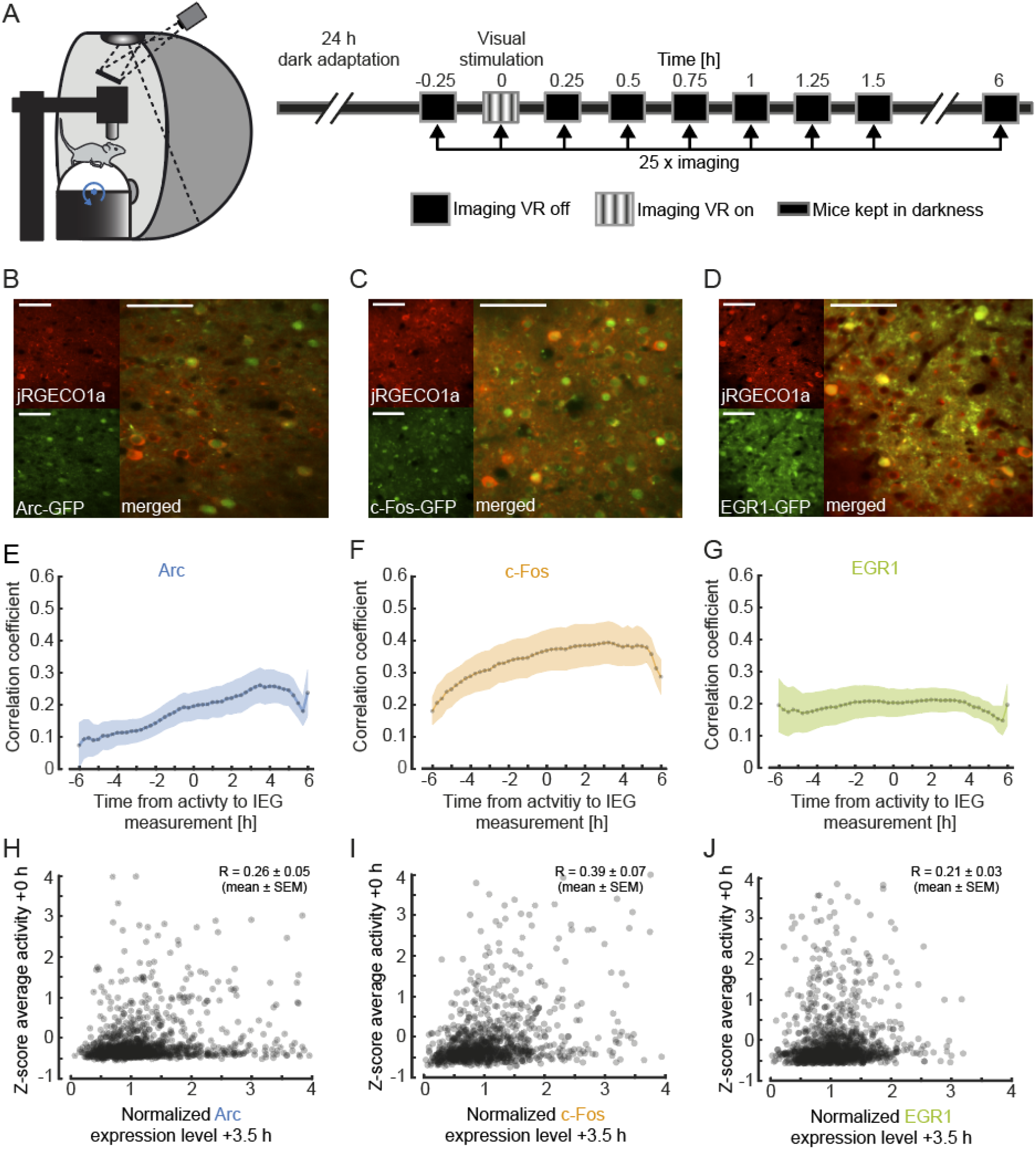
Simultaneous imaging of neuronal activity and immediate early gene expression in visual cortex. (**A**) Left: Schematic of the virtual reality setup used for imaging experiments. Right: Schematic of the experimental timeline. Mice were dark-adapted for 24 hours. Neuronal activity and IEG expression levels were recorded in 25 imaging sessions starting immediately before and continuing until 6 hours after visual stimulation in intervals of 15 minutes. (**B**) Example two-photon images of neurons in primary visual cortex labelled with jRGECO1a (red, top left), Arc (green, bottom left), and the overlay (right). Scale bar is 50 ⍰m. (**C**) Same as in (**B**), but for c-Fos. (**D**) Same as in (**B**), but for EGR1. (**E**) Correlation of average activity and IEG expression level as a function of the time difference between the two measurements. Positive values correspond to activity measurement that preceded the IEG measurement. Dotted line indicates average correlation, shading indicates standard error of the mean (SEM) across mice (n = 4). (**F**) Same as in (**E**), but for c-Fos mice (n = 4). (**G**) Same as in (**E**), but for EGR1 mice (n = 4). (**H**) Scatter plot of Arc expression 3.5 hours after visual stimulation and average neuronal activity during visual stimulation (1382 neurons in 4 mice, 83 neurons outside of plot range). Shown in the panel is the average correlation coefficient across mice (mean ± SEM, n= 4). (**I**) Same as in (**H**), but for c-Fos (1070 neurons in 4 mice, 28 neurons outside of plot range). (**J**) Same as (**H**), but for EGR1 (1319 neurons in 4 mice, 18 neurons outside of plot range).

In visual cortex, both first visual and first visuomotor exposure are associated with significant changes in functional responses (Attinger et al., 2017). To investigate whether visual and visuomotor exposure are also associated with differential expression of IEGs, we proceeded to apply the same methods of measuring IEG dynamics and neuronal calcium activity during a mouse’s first exposure to visual input and subsequent first exposure to normal visuomotor coupling. We reared mice in complete darkness and quantified both IEG expression levels and neuronal activity before and after mice were exposed to visual input for the first time in life as well as during a subsequent phase of visuomotor learning. Under normal conditions, first visual exposure is coincident with exposure to normal visuomotor coupling. At eye opening, mice are capable of moving eyes, head, and body and thus immediately experience self-generated visual feedback. To experimentally separate the moment of first visual exposure from first exposure to normal visuomotor coupling, we recorded neuronal activity and IEG expression as mice transitioned through three different experimental conditions. Prior to experiments, three groups of mice were reared in complete darkness until postnatal day 40 (7 EGFP-Arc mice, 5 c-Fos-GFP mice, and 4 EGR1-GFP mice). We then imaged neuronal activity and IEG expression levels every 12 hours for a total of 6 days. During all two-photon imaging experiments, mice were head-fixed on a spherical treadmill. During the first four recording sessions, mice were kept on the setup in darkness to measure locomotion-related and non-visual activity and remained dark housed in between recording sessions (condition 1). At the beginning of the 5^th^ recording session, mice were then exposed to visual input for the first time in their life. In the subsequent four recording sessions, mice were exposed to different virtual environments but remained housed in darkness in the time between the recording sessions (condition 2). In addition to recording activity in darkness, recording sessions in condition 2 also contained 8 min of closed-loop feedback during which visual flow on the walls of a virtual corridor was coupled to the mouse’s locomotion on the spherical treadmill. During closed-loop feedback, we added brief halts of visual flow to probe for visuomotor mismatch responses (Keller et al., 2012). This was followed by a phase of open-loop feedback during which the visual flow generated by the mouse during the closed-loop feedback was replayed independently of the locomotion of the mouse. Lastly, we presented a series of drifting gratings to the mouse to quantify visual responses (see Methods). Following recording session 8, mice were introduced to a normal 12 h light / 12 h dark cycle. At this time, mice first experienced normal visuomotor coupling in their home cage. We continued recording for an additional four sessions (condition 3) with the same series of closed-loop, open-loop and grating stimulation phases as in condition 2 (**Figure 2A**). Recording sessions lasted on average 12 min ± 0.5 min (mean ± SEM) in condition 1, and 83 min ± 1 min (mean ±SEM) in conditions 2 and 3 (**Figure S2**). Note, to probe for different visuomotor mismatch responses, condition 2 already contains a short period of head-fixed, closed-loop feedback. Given the short duration and the fact that coupling here is only in a small subspace (forward locomotion coupled to backward visual flow and normal coupling for eye movements) of the total space of visuomotor coupling, we would expect the exposure to normal visuomotor coupling in condition 3 to have a stronger influence on visuomotor integration. The aim of this paradigm was to experimentally separate the first visual exposure at the beginning of condition 2, from the first exposure to normal visuomotor coupling at the beginning of condition 3.

**Figure 2.**
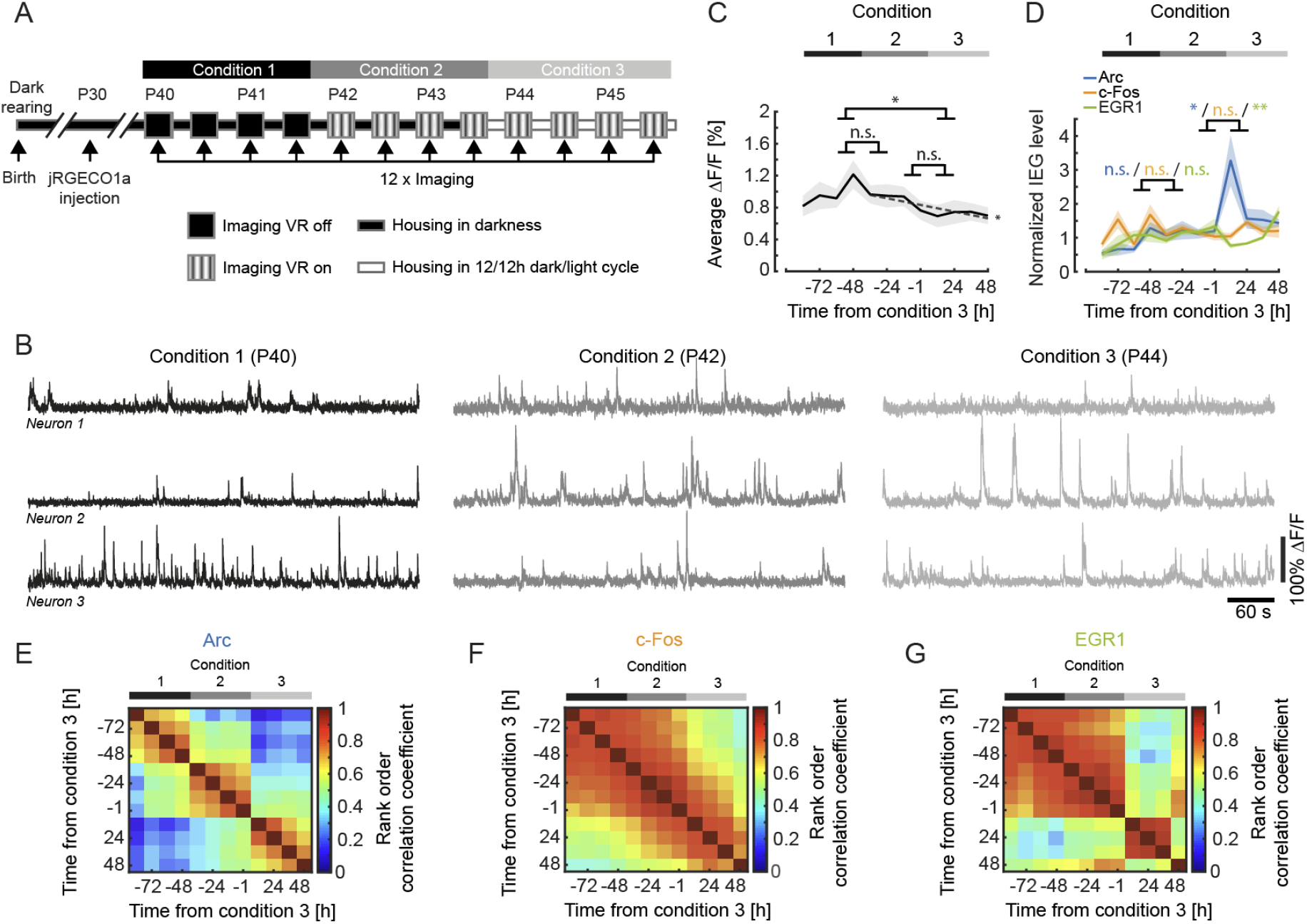
IEG expression dynamics during visuomotor learning. (**A**) Schematic of the experimental timeline. Mice were born and reared in complete darkness. jRGECO1a was injected 10 to 12 days prior to the start of imaging experiments. We then imaged calcium activity and IEG expression levels every 12 h over the course of 6 days both before and after first visual exposure and first exposure to normal visuomotor coupling. On the first two days (condition 1) activity in visual cortex was recorded in complete darkness while mice were head-fixed and free to run on a spherical treadmill. On the third day of recording mice were exposed to visual feedback (first visual exposure) in a virtual reality environment. Outside of the recording sessions mice were still housed in complete darkness (condition 2). Starting on day 5, mice were subjected to a 12 h / 12 h light/dark cycle (condition 3). (**B**) Example calcium activity of 3 neurons recorded in the same mouse in the dark across all three conditions. (**C**) Average calcium activity during all conditions (condition 1 vs. 2: p = 0.2183, condition 2 vs. 3: p = 0.527, condition 1 vs. 3: p = 0.0123, 5067 neurons, paired t-test). Shading is SEM over mice. Dashed line indicates linear fit to the data of conditions 2 and 3. The linear fit to the data from conditions 2 and 3 exhibited a significant negative slope (p = 0.0293, R^2^ = 0.371, linear trend analysis, see Methods). (**D**) Normalized mean IEG expression levels during all conditions. Expression level of Arc (blue, 1969 neurons in 7 mice) significantly increased after first exposure to visuomotor coupling, decreased for EGR1 (green, 1213 neurons in 4 mice) and remained unchanged for c-Fos (orange, 1885 neurons in 5 mice). Change in IEG expression level between conditions 1 and 2 for Arc: 0.1764 ± 0.1556, p = 0.2775; c-Fos: −0.0536 ± 0.1877, p = 0.7816; EGR1: −0.0371 ± 0.1246, p = 0.7745 (mean ± SEM, paired t-test). Change in IEG expression level between conditions 2 and 3 for Arc: 1.2628 ± 0.5012, p = 0.0256; c-Fos: 0.01612 ± 0.1372, p = 0.2702; EGR1: −0.4568 ± 0.1130, p = 0.0049 (mean ± SEM, t-test). Shading indicates SEM over mice. (**E**) Average rank order correlation coefficients for Arc expression during visuomotor learning (7 mice). The expression pattern changes both at the onset of conditions 2 and 3. (**F**) Same as in (**E**), but for c-Fos (5 mice). The expression pattern exhibits no apparent transitions. (**G**) Same as in (**E**), but for EGR1 (4 mice). The expression pattern changes at the onset of condition 3.

It has been shown that visual input can drive the expression of different IEGs in a subset of neurons in visual cortex (Kaminska et al., 1996; Kawashima et al., 2013; Rosen et al., 1992; Tagawa et al., 2005; Wang et al., 2006). Based on this it is sometimes assumed that neuronal activity in visual cortex is higher with visual input than it is in darkness. Differences in neuronal activity between light and dark, however, are small and predominantly transient (Fiser et al., 2004; Hengen et al., 2016; Torrado Pacheco et al., 2019). To test whether in our paradigm first visual exposure or first exposure to normal visuomotor coupling results in an increase of average neuronal activity in visual cortex, we quantified average neuronal activity in each recording session (in condition 1 this only included recordings in darkness, while in conditions 2 and 3 this included recordings in darkness, closed and open-loop feedback, as well as drifting gratings). Consistent with a strong motor-related drive in visual cortex (Keller et al., 2012; Saleem et al., 2013) and rapid homeostatic restoration of average activity following removal of visual input (Keck et al., 2013), we found no evidence of an increase of average neuronal activity at the onset of either condition 2 (first visual exposure) or condition 3 (first exposure to normal visuomotor coupling) (**Figures 2B and 2C**). To the contrary, following the first visual exposure, there was a trend for decreasing activity levels (p = 0.0293, R^2^ = 0.371, linear trend analysis, see Methods). We next quantified average expression of Arc, c-Fos and EGR1 over the same time course. Consistent with the absence of a change in average activity levels, we found no significant changes in the expression levels of any of the three IEGs following the first visual exposure at the beginning of condition 2 (**Figure 2D**). Note, we cannot exclude that there is a transient increase in IEG expression between 1 h and 12 h following first visual exposure, as we only recorded for 1 hour every 12 hours. We did however find that the first exposure to normal visuomotor coupling at the beginning of condition 3, resulted in an increase in the expression of Arc and a decrease in the expression of EGR1 in the absence of a measurable change in average neuronal activity levels (**Figure 2D**). To test for changes in the pattern of IEG expression, we quantified the similarity of IEG expression patterns by computing the correlation of IEG expression vectors between imaging time points (see Methods). We found that the pattern of Arc expression changed both with the first visual exposure (onset of condition 2) and the first exposure to normal visuomotor coupling (onset of condition 3) (**Figure 2E**). The pattern of c-Fos expression exhibited no detectable discontinuous changes (**Figure 2F**), while the pattern of EGR1 expression exhibited a marked transition with the first exposure to normal visuomotor coupling (onset of condition 3) (**Figure 2G**). This suggests that the expression patterns of IEGs are differentially and dynamically regulated by visuomotor experience, also in absence of population mean expression level changes.

Neurons in layer 2/3 of primary visual cortex are driven differentially by visual and motor-related inputs (Attinger et al., 2017; Keller et al., 2012; Leinweber et al., 2017). Given that the expression patterns of the three IEGs are differentially altered by first visual exposure and first exposure to normal visuomotor coupling, we speculated that the different IEGs could be preferentially expressed in different functional types of excitatory neurons in layer 2/3. Neurons that are more strongly visually driven, likely by bottom-up visual input, could have a different IEG expression profile than neurons that are more strongly driven by top-down motor-related signals (Leinweber et al., 2017; Makino and Komiyama, 2015). To test this, we quantified the functional properties of the neurons with the highest IEG expression levels immediately after the first exposure to normal visuomotor coupling where we observed the largest mean IEG expression level changes (**Figure 2D**). We selected the 10 % of neurons with the highest Arc, c-Fos and EGR1 expression, respectively, at the beginning of condition 3 (Arc: 197 neurons, c-Fos: 189 neurons, EGR1: 121 neurons) and tested whether these neurons were more strongly driven by visual or motor-related input. As a measure of the strength of the motor-related input, we used the magnitude of the neuronal response during running onsets in darkness. We found that on average neurons with high EGR1 expression levels developed higher motor-related responses than the rest of the population in both condition 2 and condition 3. Conversely, neurons with high Arc expression levels on average developed motor-related responses that are lower than the rest of the population following exposure to normal visuomotor coupling. Responses in neurons with high c-Fos expression levels were not different from responses in the rest of the population (**Figure 3A**). To quantify the strength of visual input we used the magnitude of the neuronal response to drifting grating stimuli. Consistent with the fact that Arc expression can be selectively induced by visual stimuli in a stimulus-specific manner (Kawashima et al., 2013), we found that neurons with high Arc expression levels developed responses to drifting grating stimuli that were on average stronger than the rest of the population after exposure to normal visuomotor coupling. The drifting grating responses of neurons with high EGR1 or c-Fos expression levels were not different from the mean population response (**Figure 3B**). Thus, neurons with high levels of EGR1 expression after first exposure to normal visuomotor coupling were more strongly driven by motor-related input, while those with high levels of Arc expression were more strongly driven by visual input.

**Figure 3.**
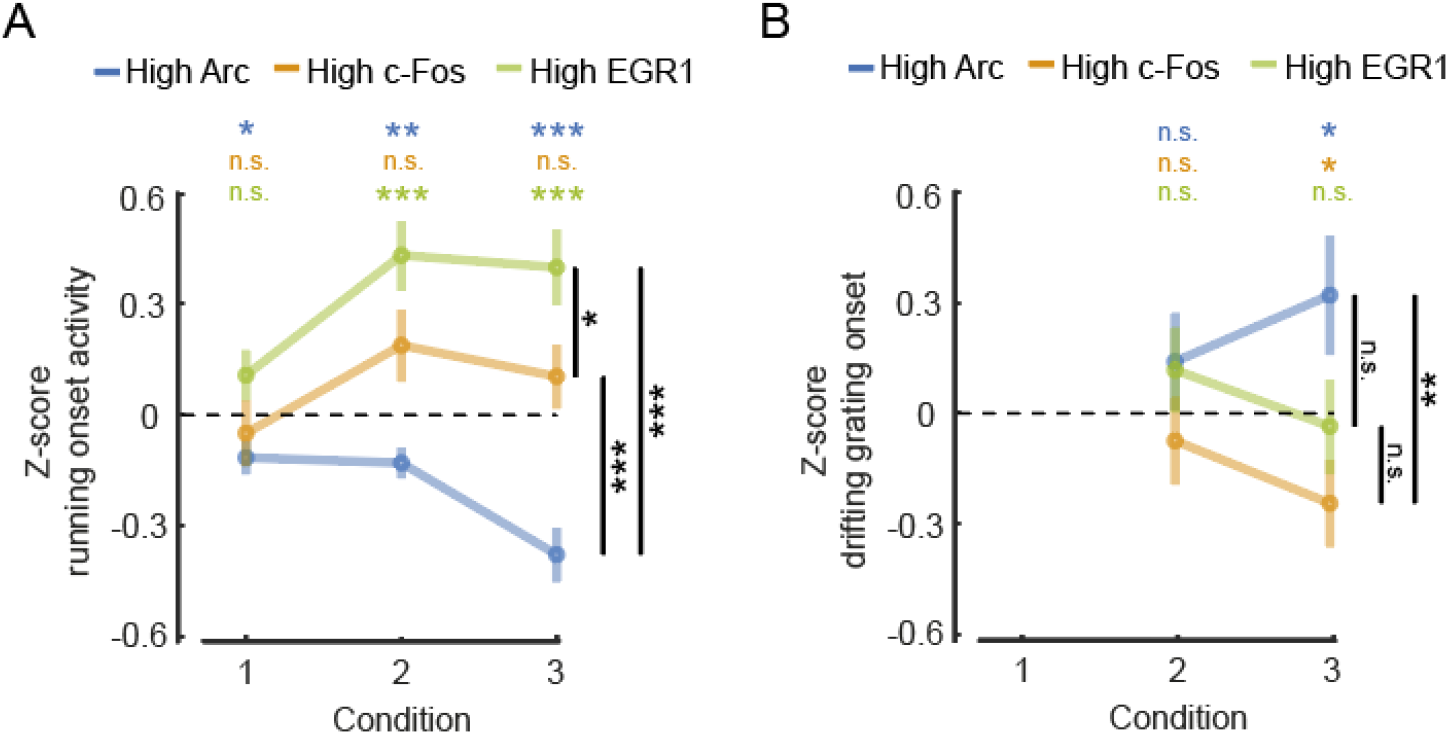
Differential relationship between IEG expression and motor-related and visual responses. (**A**) Average running onset response during darkness for the top 10 % IEG expressing neurons (Arc: 197 neurons, c-Fos: 189 neurons, EGR1: 121 neurons). Neuronal responses were pooled from all experimental sessions for each condition, subtracted by the mean and normalized by the standard deviation of the response of all neurons (Z-score). Error bars are SEM over neurons. Statistics above the plot indicate comparisons against 0, statistics to the right are between-group comparisons. n.s.: p > 0.05, *: p < 0.05, **: p < 0.01, *** p < 0.001, t-test. (**B**) Average grating onset response for the top 10 % IEG expressing neurons (Arc: n = 197, c-Fos: n = 189, EGR1: n = 121). Neuronal responses were pooled from all experimental sessions for conditions 2 and 3, subtracted by the mean and normalized by standard deviation of the response of all neurons (Z-score). Error bars are SEM over neurons. Statistics above the plot indicate comparisons against 0, statistics to the right are between-group comparisons. n.s.: p > 0.05, *: p < 0.05, **: p < 0.01, *** p < 0.001, t-test.

One of the signals that has been speculated to be computed in mouse primary visual cortex that combines visual and motor-related input is visuomotor mismatch (Attinger et al., 2017; Keller et al., 2012; Zmarz and Keller, 2016). Neurons that respond to mismatch, or negative prediction errors, are thought to receive excitatory motor-related input and inhibitory visual input (Attinger et al., 2017; Keller and Mrsic-Flogel, 2018). We speculated that given the increased motor-related activity in neurons that express high levels of EGR1, neuronal activity in these neurons should correlate positively with running, while activity in neurons that express high levels of Arc should correlate positively with visual flow. To quantify this, we computed the correlation of neuronal activity with either running or visual flow during the open-loop phases in conditions 2 and 3 for the three groups of neurons with high IEG expression levels. We found that the activity of neurons expressing high levels of EGR1 correlated most strongly with running, while the activity of neurons with high levels of Arc expression correlated positively with visual flow (**Figure 4A**). Consistent with this we found that visuomotor mismatch responses were larger in neurons with high EGR1 expression than in the rest of the population, while they were lower in neurons with high Arc expression than in the rest of the population (**Figure 4B**). This indicates that, at the onset of normal visuomotor coupling, EGR1 is preferentially expressed in mismatch neurons or, more generally, in neurons that are driven by excitatory top-down input, while Arc is preferentially expressed in neurons that are driven by bottom-up visual input.

**Figure 4.**
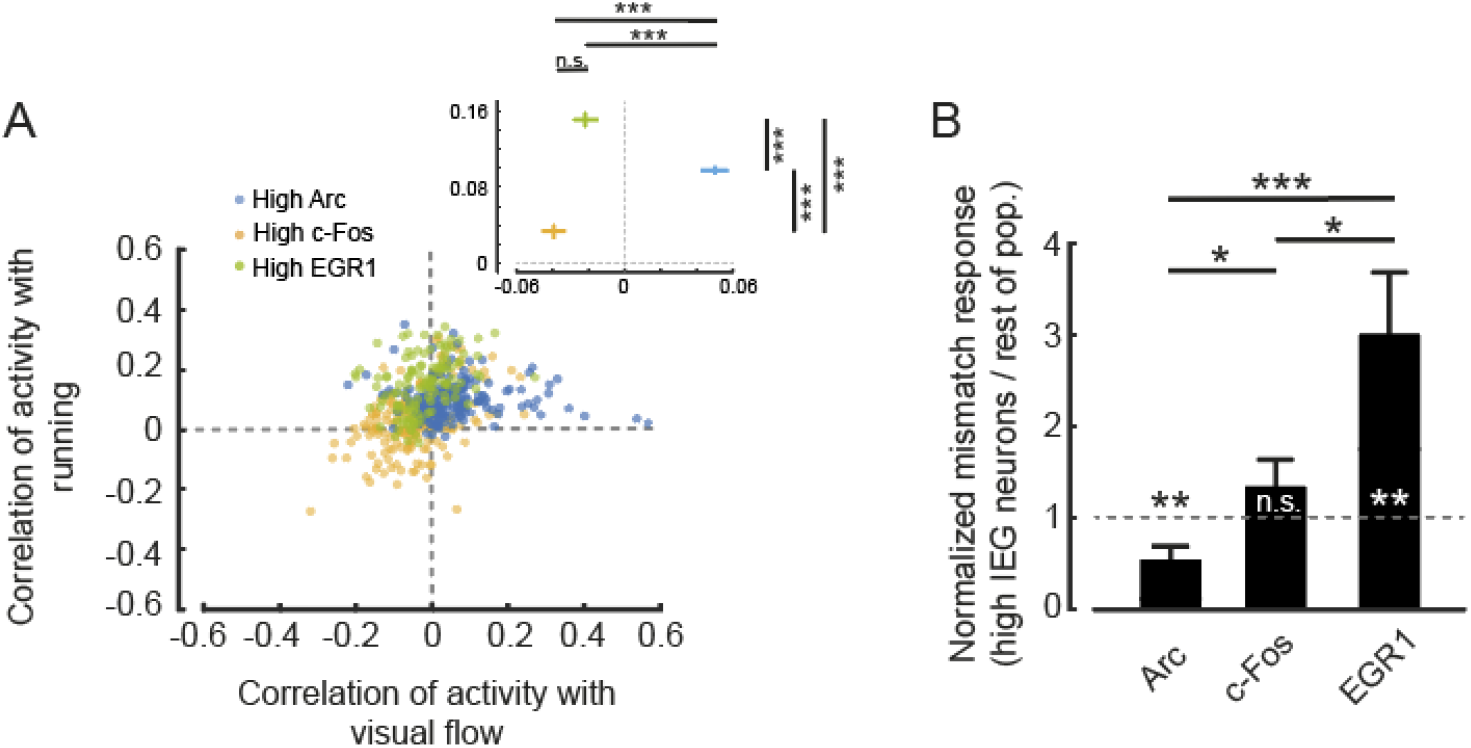
Functional cell type specific expression of IEGs in visual cortex. (**A**) Correlation of neuronal activity with running and of neuronal activity with visual flow during open-loop phases of conditions 2 and 3 for the top 10 % IEG expressing neurons (Arc: 197 neurons, c-Fos: 189 neurons, EGR1: 121 neurons). Inset: Average correlation coefficient for the three groups of high IEG expressing neurons. High Arc expressing neurons had the highest correlation with visual flow (Arc vs. c-Fos: p < 10^-10^, Arc vs. EGR1: p < 10^-9^, c-Fos vs. EGR1: p = 0.0791, t-test), while high EGR1 expressing neurons had the highest correlation with running (Arc vs. c-Fos: p < 10^-10^, Arc vs. EGR1: p < 10^-8^, c-Fos vs. EGR1: p < 10^-10^, t-test). (**B**) Mismatch responses in condition 3 were significantly higher for the top 10 % EGR1 expressing neurons and significantly lower for the top 10 % Arc expressing neurons than the rest of the respective population (Arc: 197 neurons, c-Fos: 189 neurons, EGR1: 121 neurons). Arc: p = 0.0461, c-Fos: p = 0.2273, EGR1: p = 0.0234; Arc vs. c-Fos: p = 0.0101, c-Fos vs. EGR1: p = 0.048, Arc vs. EGR1: p = 0.0057, t-test.

## Discussion

It is well established that both neuronal activity and plasticity are linked to the expression of immediate early genes (Dudek, 2008; Mahringer et al., 2019; Minatohara et al., 2015; Wang et al., 2021; Yap and Greenberg, 2018). Comparably little, however, is known about how specific functional characteristics of neurons relate to the expression of immediate early genes. Here we investigated the relationship between the expression of three IEGs (Arc, c-Fos, and EGR1) and functional responses in excitatory layer 2/3 neurons of mouse visual cortex. We found that during visuomotor learning following a mouse’s first visual exposure in life, Arc was preferentially expressed in neurons that are driven by excitatory bottom-up visual input, while EGR1 was preferentially expressed in neurons that are driven by motor-related input. In addition, we found that neurons expressing high levels of EGR1 exhibit visuomotor mismatch responses higher than the rest of the population, while neurons expressing high levels of Arc exhibit visuomotor mismatch responses weaker than the rest of the population.

Such a relationship between a neuron’s IEG expression profile and its functional properties could be explained by differences in the contribution of different IEGs to different types of input synapses. Arc, c-Fos, and EGR1 all have unique cellular functions, and it is conceivable that they make different contributions to different synapse types. Genes for a subset of GABA_A_ receptor subunits, for example, are transcriptional targets of EGR1 (Mo et al., 2015). If the postsynaptic subunit composition of the GABA receptor is correlated with the presynaptic inhibitory cell type, EGR1 expression could preferentially upregulate specific inhibitory input pathways. Similar input pathway-specific roles have been described for other IEGs. Neuronal activity-regulated pentraxin (NARP) is secreted by pyramidal neurons and exclusively accumulates at parvalbumin-positive inhibitory neurons where it regulates excitatory synapses onto these cells (Chang et al., 2010; Gu et al., 2013). The activity-dependent transcription factor NPAS4 has been found to restrict the number of synapses of mossy-fiber input specifically onto CA3 pyramidal cells during learning (Weng et al., 2018).

Our data would be consistent with the interpretation that the IEG expression pattern of a given neuron correlates with its pattern of synaptic inputs. Neurons that predominantly receive excitatory bottom-up drive likely require a different distribution and type of input synapses compared to neurons that receive mainly top-down excitatory drive. Layer 2/3 neurons that exhibit strong motor-related and mismatch responses are thought to be driven by top-down excitatory inputs (Leinweber et al., 2017), which predominantly target apical dendrites (Petreanu et al., 2009). Conversely, layer 2/3 neurons with strong visual responses are thought to be driven by bottom-up visual inputs, which predominantly target basal dendrites (Petreanu et al., 2009). We have speculated that mismatch neurons that receive motor-related input also receive matched bottom-up inhibitory input from a specific subset of somatostatin (SST)-positive interneurons (Attinger et al., 2017). Thus, EGR1 expression may be preferentially increased in neurons that are driven by excitatory top-down input and SST mediated bottom-up inhibition, while Arc expression may be preferentially increased in neurons that are driven by excitatory bottom-up visual input. This may explain why a change to the visual input alone at first visual exposure primarily resulted in a rearrangement of the Arc expression pattern (**Figure 2E**) but left the EGR1 expression pattern relatively unaffected (**Figure 2G**), while first exposure to normal visuomotor coupling resulted in a rearrangement of the expression pattern of both Arc and EGR1. It is also consistent with the fact that experience dependent changes of EGR1 expression are primarily observed in superficial layer 2/3 neurons (Xie et al., 2014) that likely are more strongly targeted by top-down input than deep layer 2/3 neurons.

When interpreting our results, it should be kept in mind that both the method we use to approximate IEG expression levels and the method we use to approximate neuronal activity levels come with a series of caveats. In the case of the transgenic mice used for the IEG expression measurements, two of these express a fusion protein (Arc and c-Fos), where the IEG is likely overexpressed (Steward et al., 2017), and it is possible that the decay kinetics of the fusion protein differ from those of the native protein. In the case of the GFP driven by the *egr1* promoter, the GFP decay kinetics are likely different from the decay kinetics of EGR1. However, these potential differences in decay kinetics and expression levels do not completely mask the correlation between IEG expression levels and reporter proteins. In post-mortem histological stainings the expression of GFP in these mouse lines overlaps well with the expression levels of the IEGs (Barth et al., 2004; Okuno et al., 2012; Wang et al., 2021; Xie et al., 2014; Yassin et al., 2010). Thus, reporter protein levels reflect a filtered version of IEG expression levels, and the two are likely related by a monotonic function. Given that all our analyses rely only on relative expression levels among populations of simultaneously recorded neurons or relative changes of expression levels in time, the lack of a direct measurement of IEG expression levels should not change our conclusions. A second caveat concerns the genetically encoded calcium indicator used to measure neuronal activity. Our activity measures are biased towards bursts of neuronal activity, as single spikes are probably not always detectable using calcium indicators *in vivo.* However, even though the transfer function from neuronal activity to calcium signal is non-linear, it is monotonic. Thus, we may be underestimating the correlation between neuronal activity and IEG expression, but neither caveat would bias the results towards finding specific correlations between different IEGs and functional cell types. Lastly, to experimentally separate first visual exposure from first visuomotor exposure, and to be able to record neuronal activity throughout this paradigm, we had to dark rear mice from birth. Dark rearing is known to delay normal development of V1 (Hensch, 2005; Sherman and Spear, 1982). However, using a similar experimental paradigm, we have previously found that dark rearing did not impair development of visuomotor integration once mice are exposed to normal visuomotor coupling (Attinger et al., 2017). Thus, given that our conclusions are based on differences between different groups of mice that were all dark reared, the caveats associated with the dark rearing are unlikely to substantially alter our conclusions.

In summary, our results suggest that the expression of Arc and EGR1 in layer 2/3 neurons in mouse visual cortex may be a correlate of the type of functional input the neuron receives. Such a preference for expression in a functionally specific subset of neurons would be consistent with differential changes in the ratio of the expression of different IEGs under conditions that result in identical mean levels of neuronal activity (Bailey and Wade, 2003; Farina and Commins, 2016; Guzowski et al., 2006) that are difficult to explain if IEG expression were simply driven by mean activity. In future experiments, it will be important to establish a more detailed picture of how immediate early genes could orchestrate or stabilize the pattern of functionally distinct input streams a neuron receives.

## Methods

### Animals and surgery

All animal procedures were approved by and carried out in accordance with guidelines of the Veterinary Department of the Canton Basel-Stadt, Switzerland. We used imaging data from a total of 11 EGFP-Arc mice (Okuno et al., 2012), 9 c-Fos-GFP mice (Barth et al., 2004) and 8 EGR1-GFP mice (Xie et al., 2014), aged 40 days at the start of visuomotor learning (**Figures 2 – 4**) or aged 100-104 (Arc), 279-291 (c-Fos) or 120-124 (EGR1) days (**Figure 1**). Sample sizes were chosen according to the standards in the field and no statistical methods were used to predetermine sample sizes. Mice were group-housed in a dark cabinet and in a vivarium (light/dark cycle: 12 h / 12 h). Viral injections and window implantation were performed as previously described (Dombeck et al., 2010; Leinweber et al., 2014). Briefly, for sensorimotor learning experiments, mice (aged 29 d ± 1 d, mean ± SEM) were anesthetized in darkness using a mix of fentanyl (0.05 mg/kg), medetomidine (0.5 mg/kg) and midazolam (5 mg/kg), and additionally their eyes were covered with a thick, black cotton fabric during all surgical procedures. A 3 mm to 5 mm craniotomy was made above visual cortex (2.5 mm lateral of lambda (Paxinos and Franklin, 2013)) and AAV2/1-Ef1a-NES-jRGECO1a-WPRE ((Dana et al., 2016); titer: between 7.2×10^10^ GC/ml and 6.8 × 10^12^ GC/ml) was injected into the target region. The craniotomy was sealed with a fitting cover slip. A titanium head bar was attached to the skull and stabilized with dental cement.

### Imaging and virtual reality

Imaging commenced 10 – 12 (visuomotor learning experiments, **Figures 2 – 4**) or 12 – 29 (**Figure 1**) days following virus injection and was carried out using a custom-built two-photon microscope. Illumination source was a Chameleon Vision laser (Coherent) tuned to a wavelength of either 950 nm, 990 nm or 1030 nm. Imaging was performed using an 8 kHz resonance scanner (Cambridge Technology) resulting in frame rates of 40 Hz at a resolution of 400 × 750 pixels. In addition, we used a piezo actuator (Physik Instrumente) to move the objective (Nikon 16x, 0.8 NA) in steps of 15 μm between frames to acquire images at four different depths, thus reducing the effective frame rate to 10 Hz. The behavioral imaging setup was as previously described (Leinweber et al., 2014). After brief isoflurane anesthesia mice were head-fixed in complete darkness and the setup was light-shielded before every imaging session. Mice were free to run on an air-supported polystyrene ball, the motion of which was restricted to the forward and backward directions by a pin. The ball’s rotation was coupled to linear displacement in the virtual environment that was projected onto a toroidal screen surrounding the mouse. The screen covered a visual field of approximately 240 degrees horizontally and 100 degrees vertically. All displayed elements of the tunnel or sinusoidal gratings were calibrated to be isoluminant.

### Experimental design

For experiments shown in **Figure 1**, mice were dark-adapted for 24 h and 17 min ± 10 min (mean ± SEM, 12 mice) before head fixation under the microscope in darkness. Activity and immediate early gene expression were recorded every 15 minutes for 6 hours. Except for the time of visual stimulation with sinusoidal gratings moving in 8 different directions (a total of 80 presentation in random order), mice were kept in complete darkness under the microscope for the duration of the entire experiment. For visuomotor learning experiments (**Figures 2 – 4**) mice were born and reared in complete darkness until P44 and then transferred to a vivarium with a 12 h /12 h light/dark cycle. Experimental sessions started on P40 and occurred twice per day, spaced 12 h apart. In condition 1, all imaging was done in complete darkness and experiments consisted of recording approximately 8 min of neuronal activity during which mice were free to run on the spherical treadmill. IEG expression level measurements were taken before and after each activity recording. In conditions 2 and 3, neuronal activity measurements consisted of 7 recordings of approximately 8 minutes each. Each recording session started with a recording in darkness, followed by a closed-loop recording. In the closed-loop recording, the movement of the mouse in a linear virtual corridor (sinusoidal vertical grating) was coupled to the locomotion of the mouse on the spherical treadmill. During the closed-loop session we included brief (1 s) halts of visual flow to induce mismatch events (Attinger et al., 2017). The subsequent two recordings were of the open-loop type and consisted of a playback of the visual flow the mouse had generated during the preceding closed-loop recording. Subsequently, mice were exposed to a second recording in darkness, followed by a visual stimulation recording. During the visual stimulation sinusoidal moving grating stimuli (2 second standing grating, 3 second drifting grating, 8 different orientations, 10 presentations of each orientation, in a randomized order) were presented. Finally, mice were exposed to a third recording in darkness. In early phases of the experiment mice were encouraged to run by applying occasional mild air puffs to the neck.

### Data analysis

Imaging data were full-frame registered using a custom-written software (Leinweber et al., 2014). Neurons were selected manually based on their mean fluorescence or maximum projection in the red channel (jRGECO1a). This biased our selection towards active neurons. Fluorescence traces were calculated as the mean pixel value in each region of interest per frame, and were then median-normalized to calculate ΔF/F. ΔF/F traces were filtered as previously described (Dombeck et al., 2007). GFP intensities were calculated as the mean pixel value in each region of interest (ROI) for mean fluorescence projections. To compensate for expression level differences between different IEG mouse lines as well as for image quality differences between different mice we normalized the GFP level measurements as follows: For each mouse, all ROI measurements were subtracted by the minimum calculated over all ROIs and timepoints, and normalized by the median over all ROIs and timepoints.

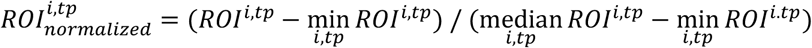

This ensured that the minimum value of IEG expression was 0 and the median 1. No blinding of experimental condition was performed in any of the analyses. Statistical tests were used as stated in the figure legends.

**Figure 1.** Examples images (**Figures 1B–1D**) are average projections of the recorded channel. IEG expression was normalized as described above (**Figures 1E–1G**). Correlation coefficients (**Figures 1E–1G**) were calculated based on the neuronal population vectors of average activity and IEG expression per measurement timepoint, for each mouse. For the statistical comparison of the correlation coefficients of IEG expression levels with neural activity between the three different groups (4 mice per group), data were bootstrapped 5 times with random replacement and then a t-test was performed on the bootstrapped data.

**Figure 2.** To compare changes in neural activity and IEG expression levels between conditions we averaged data from the last two recording sessions of the previous condition and the first two recording sessions of the following condition (**Figures 2C and 2D**). Linear trend analysis (**Figure 2C**) was performed using the MATLAB regress function treating the average activity per timepoint for each mouse as an independent observation. To quantify the significance of the linear trend we report the R^2^ statistic and p-value of the F statistic. The linear fit shown (**Figure 2C**), is the average over the linear fits performed to the data of each mouse individually using the MATLAB polyfit and polyval functions. Rank order correlation coefficients (**Figures 2E–2G**) were determined based on the population vectors of average IEG expression per measurement timepoint and mouse, and then averaged.

**Figure 3.** For plots of event-triggered activity changes DF/F traces were baseline-subtracted by the average DF/F in a window −500 ms to −100 ms preceding the event onset. Z-scores were obtained on a population vector with average stimulus onset values calculated over a response window of 1.5 s. High IEGs neurons were selected as the top 10% of IEG expressing neurons based on average expression level on the first day of condition 3.

**Figure 4.** Correlation coefficients (**Figure 4A**) were calculated by correlating each neuron’s activity trace with either the running trace or the visual flow trace during open-loop phases. High IEG neurons were selected with the same criteria used for **Figure 3**. Stimulus-triggered fluorescence changes (**Figure 4B**) were mean-subtracted in a window −500 ms to −100 ms preceding the stimulus onset. Responses were quantified in a window of 1.5 s.

**Figure S1**. Correlation coefficients of mean or maximum activity with IEG expression were calculated for each mouse (**Figure S1B**). Mice from visual stimulation experiments (**Figure 1**) and sensorimotor learning experiments (**Figures 2–4**) were pooled for this analysis. For mice from the sensorimotor learning experiments the calculation was done using mean or maximum activity of the first recording segment and the last IEG measurement within a session. Shown is the average correlation across all sessions.

## Acknowledgements

We thank the entire Keller lab for helpful discussion and comments on earlier versions of this manuscript. We thank Daniela Gerosa-Erni for production of the AAV vectors. Version 4 of this preprint has been peer-reviewed and recommended by Peer Community In Neuroscience (https://doi.org/10.24072/pci.neuro.100005).

## Author contributions

D.M. and P.Z. performed the experiments, D.M. analyzed the data. H.O. and H.B. made the EGFP-Arc mouse. All authors wrote the manuscript.

## Data, scripts, and code availability

All imaging and image processing code can be found online at https://sourceforge.net/projects/iris-scanning/ (IRIS, imaging software package) and https://sourceforge.net/p/iris-scanning/calliope/HEAD/tree (Calliope, image processing software package). All the raw data and analysis code used in this study can be downloaded from the following website: http://data.fmi.ch/PublicationSupplementRepo/.

## Supplementary Figures

**Figure S1.**
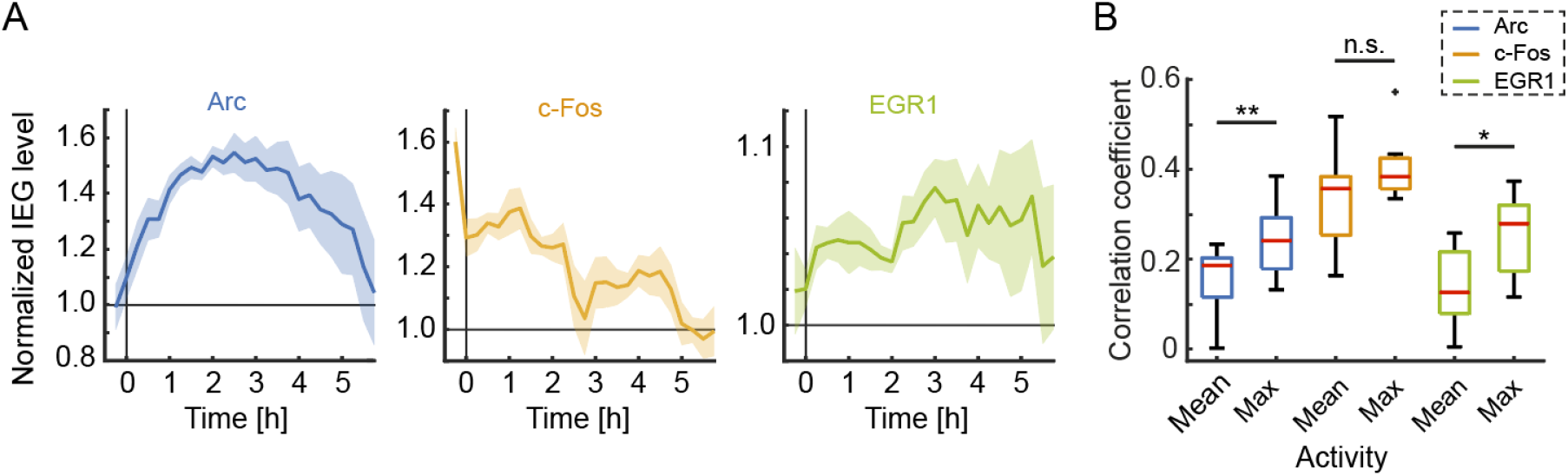
Time course of IEG expression during the imaging paradigm and correlation of IEG expression with mean and maximum neuronal activity. Related to Figure 1. (A) Time course of normalized IEG expression levels following 24 h dark adaptation and 15 min visual stimulation at time 0. Shading indicates SEM over neurons. (B) Correlation coefficient of mean and maximum activity (average across or peak within a recording session, respectively) with IEG expression 3.5 h after stimulation or recording onset (Arc: 11 mice, c-Fos: 9 mice, EGR1: 8 mice). Box whisker plot: red line indicates median, box marks 25th to 75th percentiles and whiskers extended to the next most extreme datapoint within a range of 1.5 times the interquartile distance (rank sum test, Arc: p = 0.0086, c-Fos: p = 0.1359, EGR1: p = 0.0207).

**Figure S2.**
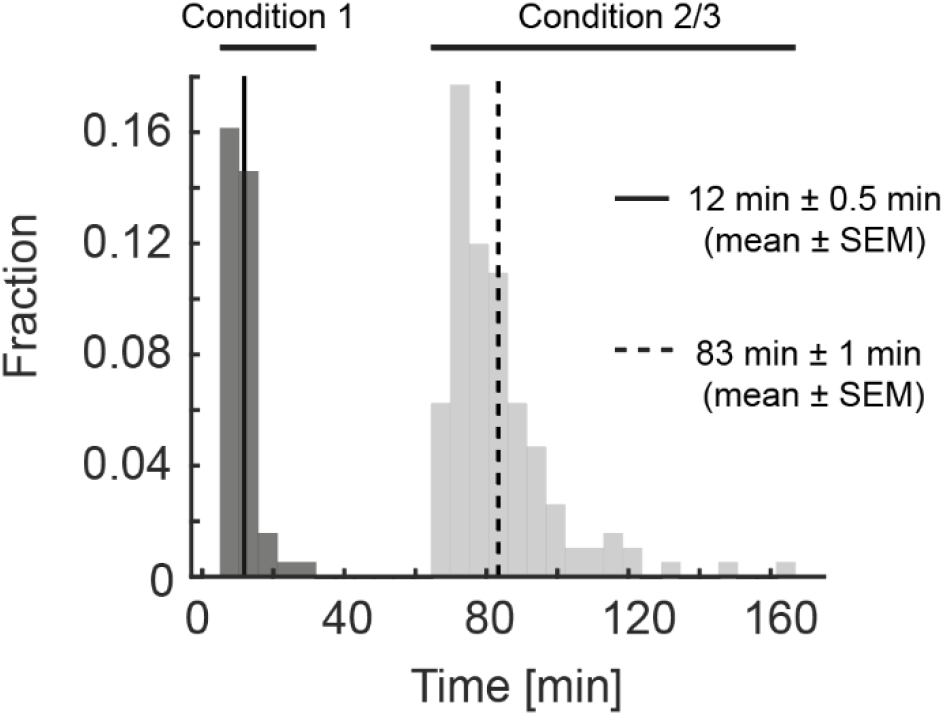
Duration of recording sessions. Related to Figure 2. Histogram of the durations of the recording sessions. On average, one recording session lasted for approximately 12 min during condition 1 (solid line) and, due to the addition of closed-loop, open-loop, and grating stimulation phases, 83 min during conditions 2 and 3 (dashed line).

